# Leveraging Transfer Learning for Predicting Protein-Small Molecule Interactions

**DOI:** 10.1101/2024.10.08.617219

**Authors:** Jian Wang, Nikolay V. Dokholyan

**Affiliations:** Department of Pharmacology, Penn State College of Medicine, Hershey, PA 17033, USA; Department of Biochemistry & Molecular Biology, Penn State College of Medicine, Hershey, PA 17033, USA; Department of Chemistry, Pennsylvania State University, University Park, PA 16802, USA; Department of Biomedical Engineering, Pennsylvania State University, University Park, PA 16802, USA

## Abstract

A complex web of intermolecular interactions defines and regulates biological processes. Understanding this web has been particularly challenging because of the sheer number of actors in biological systems: ∼10^4^ proteins in a typical human cell offer a plausible 10^8^ interactions. This number grows rapidly if we consider metabolites, drugs, nutrients, and other biological molecules. The relative strength of interactions also critically affects these biological processes. However, the small and often incomplete datasets (10^3^-10^4^ protein-ligand interactions) traditionally used for binding affinity predictions limit the ability to capture the full complexity of these interactions. To overcome this challenge, we developed Yuel 2, a novel neural network-based approach that leverages transfer learning to address the limitations of small datasets. Yuel 2 is pre-trained on a large-scale dataset to learn intricate structural features and then fine-tuned on specialized datasets like PDBbind to enhance the predictive accuracy and robustness. We show that Yuel 2 predicts multiple binding affinity metrics – Kd, Ki, and IC50 – between proteins and small molecules, offering a comprehensive representation of molecular interactions crucial for drug design and development.

## INTRODUCTION

The life of cells and organisms is defined by interactions between molecules that reside inside cells and those that are introduced, such as drugs, metabolites, nutrients, and toxins. This complex web of interactions results in a spatiotemporal organization of life processes. Understanding the basic principles underlying these interactions and being able to predict them have profound implications for biology and medicine. Yet, the fundamental physical uncertainties and the large space of interacting agents limit our abilities to fully comprehend this complex web of interactions that define biological life. A typical human cell contains approximately 10^4^ different proteins, each of which can engage in numerous interactions with other proteins, as well as with various small molecules. These interactions are essential for the proper functioning of biological systems, influencing everything from cellular signaling to metabolic pathways. However, the potential number of interactions within a single cell, potentially reaching 10^8^, poses a significant challenge to our understanding and predictive capabilities. This complexity is further compounded by the fact that these interactions vary in strength and specificity, critically affecting the stability and dynamics of biological networks. Predicting these interactions and their relative affinities remains one of the most daunting tasks in computational biology and drug discovery.

Traditional approaches to predicting protein-small molecule interactions have relied heavily on datasets with experimental protein-small molecule binding affinities, such as PDBbind^1^ and BindingDB^2^. Despite the advances in deep learning methods and their demonstrated higher accuracy across various datasets, traditional docking software such as AutoDock Vina^3^, Glide SP^4^, and MedusaDock^5,6^ remain prevalent in virtual screening. The persistent use of conventional docking methods rather than machine learning methods can be attributed to the limited size of existing datasets used to train machine learning models. These datasets typically comprise 10^3^ to 10^4^ protein-ligand interactions. For example, the PDBbind^1^ refined dataset comprises 5316 protein-ligand pairs, and the PDBbind^1^ general dataset comprises 14127 protein-ligand pairs. The small size and incomplete nature of these datasets restrict the ability of computational models to generalize across the vast interaction space. As a result, predictions made by these models may lack the accuracy and robustness needed for practical applications in drug design and other areas of biomedical research. In contrast, traditional docking software relies on physical force fields, offering a robust, theoretically based approach for predicting binding interactions. However, the accuracy of these traditional scoring functions is often constrained by their reliance on fundamental physical approximations of inter-atomic interactions.

In addition to the challenge of limited dataset size in training machine learning models, both traditional docking methods and neural networks also face difficulties in accurately predicting complex phenomena like *activity cliffs*^7,8^, where small molecular changes can drastically impact binding affinities. Encountered in complex biochemical systems, these cliffs result in the inaccurate prediction of binding affinities and limit the applicability of existing methods. Overcoming these challenges is essential for ensuring that a prediction method navigates the intricate landscape of activity cliffs to provide comprehensive and reliable insight into the dynamic interplay between proteins and bioactive compounds.

Finally, while many existing methods focus on predicting a single metric, such as the dissociation constant (Kd), a comprehensive approach that also predicts the inhibition constant (Ki), and half-maximal inhibitory concentration (IC50) is significantly more practical. Each of these metrics provides unique insights into different aspects of interaction dynamics. Kd measures the affinity of the ligand for the protein at equilibrium, Ki indicates the potency of an inhibitor, and IC50 quantifies how much of a substance is needed to inhibit a biological process by one-half. Predicting all kinds of metrics using only one method would offer a more detailed and accurate representation of the protein-ligand interaction, which is essential for drug design and development. Existing methods often fall short by addressing only one of these critical metrics or even mix them, thus limiting their applicability and precision.

Here, we propose Yuel 2, which utilizes transfer learning to address the limitations posed by small and incomplete datasets. We create a large-scale dataset by systematically docking multiple proteins with multiple ligands, resulting in a comprehensive dataset containing all possible protein-ligand pairs. This approach generates an expansive dataset that significantly surpasses the size of current datasets like PDBbind, which may only include a limited subset of possible interactions. By training Yuel 2 on this large-scale dataset, we enable the model to learn the intricate structural features of both proteins and ligands at a high level. Once the neural network has been pre-trained on this vast dataset to predict docking scores, we fine-tune the model on smaller, specialized datasets, such as PDBbind, to refine its predictive capabilities for specific binding affinities. This transfer learning strategy leverages the broader structural knowledge acquired during the initial training phase and adapts it to more focused tasks, enhancing the model’s generalizability and accuracy. In addition, the original version, Yuel^9^, relied solely on protein sequences and 2D ligand structures, addressing overfitting issues in protein-small molecule interaction predictions. In contrast, Yuel 2 integrates protein structural information, aiming to enhance predictive accuracy. Finally, Yuel 2 extends beyond the conventional aim of predicting Kd to encompass Ki and IC50 values for protein-small molecule interactions, and it also aims to address the challenge of accurately predicting activity cliffs.

## RESULTS

### Pre-training with large-scale docking dataset

Yuel 2 is composed of Yuel-SE (Structural Encoder) for pre-training and Yuel-AP (Affinity Predictor) for fine-tuning (Figure 1). In the pre-training phase, Yuel-SE processes protein pocket structures and small molecule 2D structures and then subjects the processed features to Yuel-AP, which predicts the docking scores. This approach allows Yuel-SE to learn comprehensive structural features from a large-scale dataset. During the fine-tuning phase, Yuel-AP is trained with the fixed parameters of Yuel-SE on a smaller experimental dataset to predict binding affinities. As part of the pre-training step in our transfer learning strategy, we evaluate the ability of Yuel 2 to predict docking scores generated by two docking software, MedusaDock and AutoDock Vina. To compile a comprehensive dataset for the pre-training, we begin by expanding the PDBbind dataset through systematic docking, pairing each protein with multiple ligands to create a large and diverse set of protein-ligand interactions. For each of these pairs, MedusaDock and AutoDock are employed to compute binding scores. MedusaScore, the scoring function of MedusaDock, includes various energy components such as VDW_A (Van der Waals attraction), VDW_R (Van der Waals repulsion), SOLV (solvation), SB (sidechain-backbone), HB_BB (hydrogen bond between backbone and backbone), HB_SB (hydrogen bond between sidechain and backbone), and HB_SS (hydrogen bond between sidechain and sidechain). Similarly, AutoDock Vina scores are composed of energy terms like gauss 1, gauss 2, repulsion, hydrophobic, and hydrogen bonding energies. Yuel 2 is pre-trained on this extensive dataset to predict these energy terms, demonstrating high accuracy. Specifically, Yuel 2 achieves a correlation coefficient of 0.97 for predicting VDW_A **(Figure 2**), a major component of MedusaScore (**Figure S1**), and a correlation coefficient of 0.94 for predicting the total MedusaScore (**Figure S2**). Yuel 2 also showed strong predictive performance for other energy components (**Figure 2**) and achieved an accuracy of 0.8 in predicting AutoDock Vina docking scores (**Figure 2**). These findings emphasize the capability of Yuel 2 capability to replicate and even enhance the predictive performance of traditional docking scores, underscoring its potential as a powerful tool for virtual screening.

**Figure 1.**
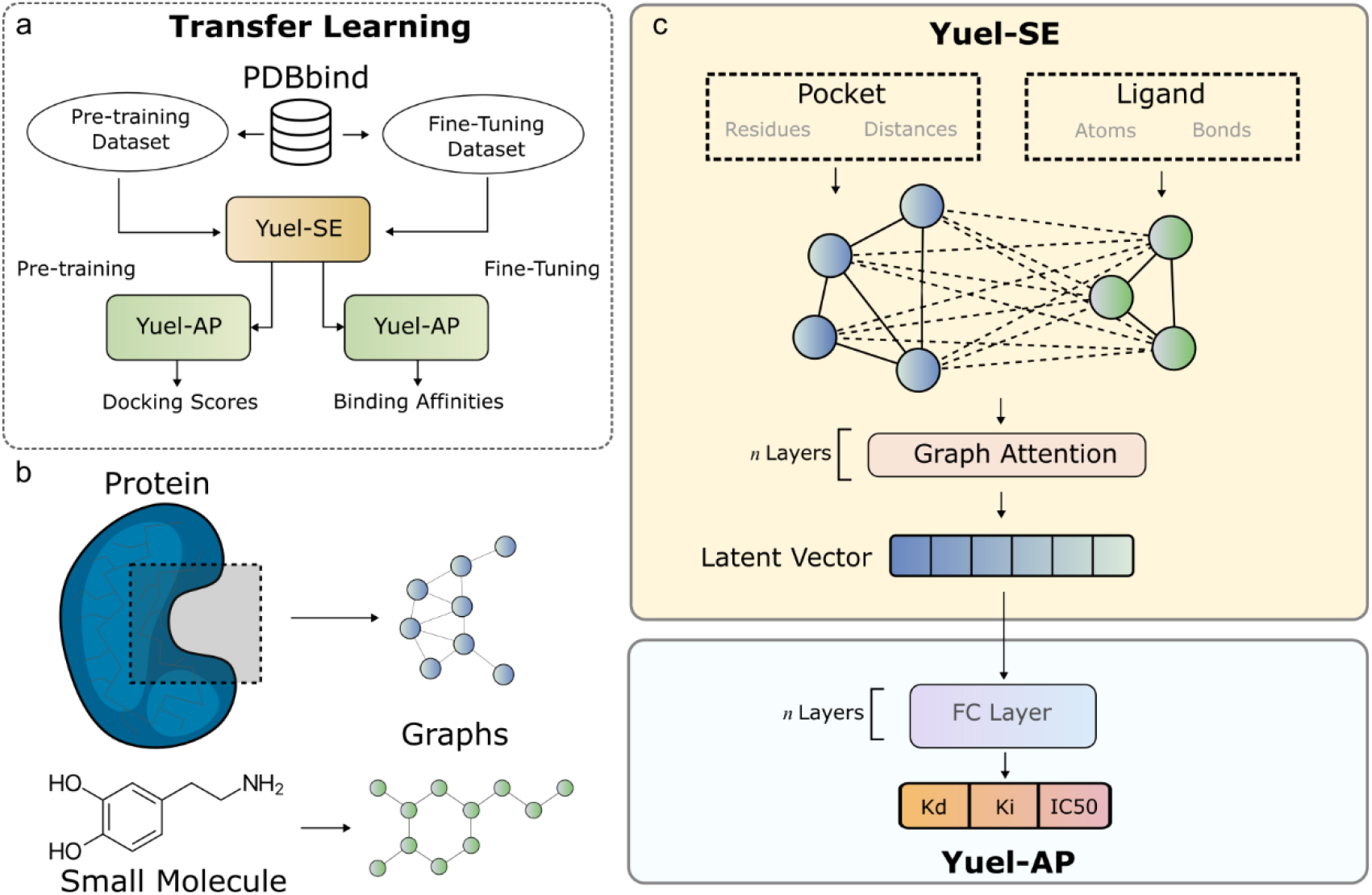
The architecture of Yuel 2. (a) We adopt transfer learning to train Yuel 2. Yuel is composed of a structure encoder (Yuel-SE) and an affinity predictor (Yuel-AP). We compile the original PDBbind dataset to a pre-training dataset and a fine-tuning dataset (Methods). Yuel 2 is first trained on the pre-training dataset, and then Yuel-SE parameters will be kept fixed, and the parameters of Yuel-AP will be re-trained on the fine-tuning dataset. (b) Proteins and small molecules are both converted to graphs. Each residue in the protein and each atom in a small molecule are represented by nodes. Residue contacts in the protein and atom bonds in the small molecule are represented by edges in graphs. (c) Yuel-SE first encodes protein pocket structure and small molecule structure to graphs. The two graphs will be merged into a large graph, which will then be subject to several graph attention layers. The latent vector generated by Yuel-SE will be subject to Yuel-AP, which is essentially several fully-connected layers.

**Figure 2.**
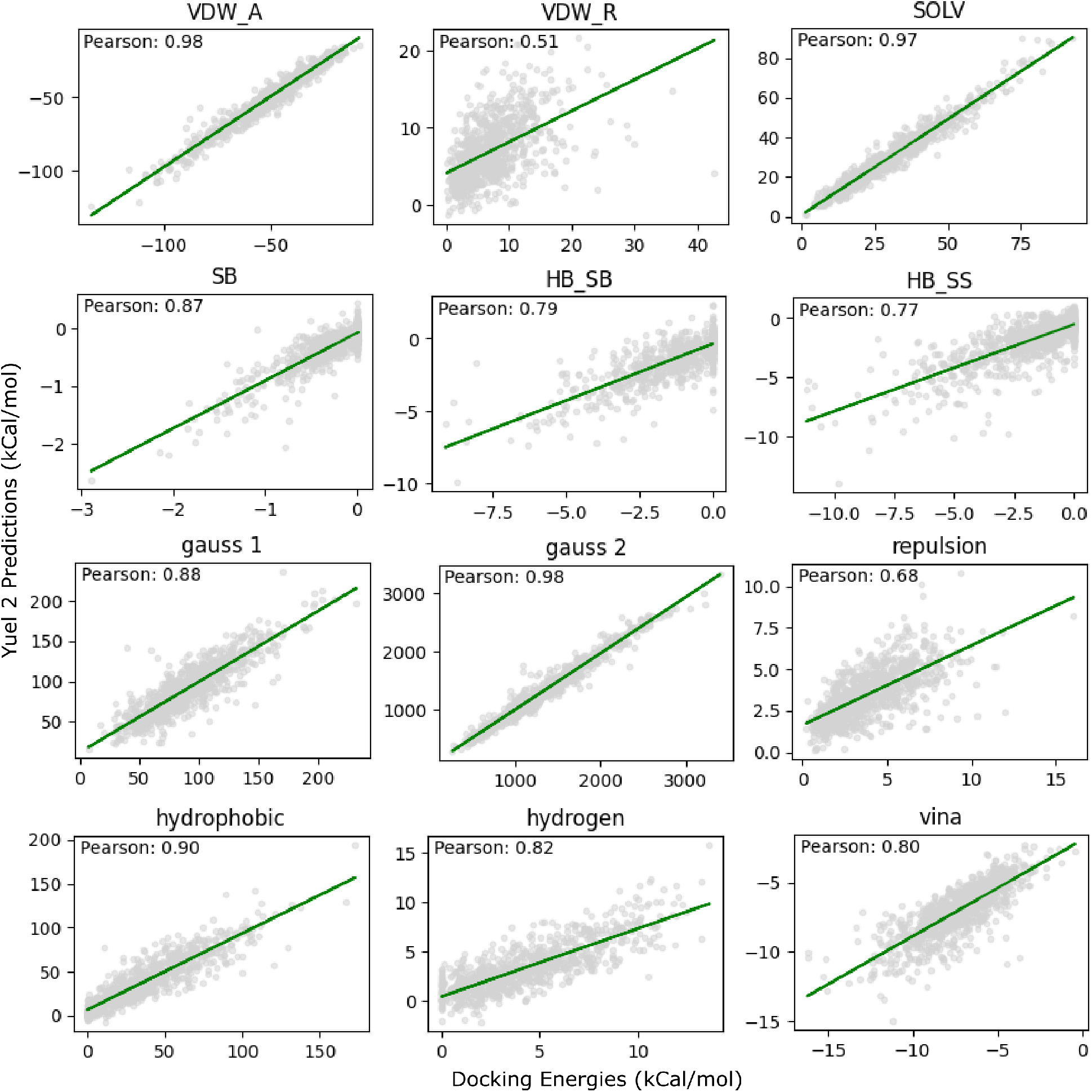
Yuel 2 recapitulates the energy items in MedusaScore and AutoDock energy. MedusaScore, the scoring function of MedusaDock, includes various energy components such as VDW_A (Van der Waals attraction), VDW_R (Van der Waals repulsion), SOLV (solvation), SB (sidechain-backbone), HB_BB (hydrogen bond between backbone and backbone), HB_SB (hydrogen bond between sidechain and backbone), and HB_SS (hydrogen bond between sidechain and sidechain). AutoDock Vina scores are composed of energy terms like gauss 1, gauss 2, repulsion, hydrophobic, and hydrogen bonding energies. Yuel 2 is pre-trained on this extensive dataset to predict these energy terms, demonstrating high accuracy.

### Fine-tuning for protein−small molecule binding affinity prediction

Using the CASF-2016^10^ test set, we evaluate the scoring power of Yuel 2 by computing the binding affinity of a protein-ligand complex based on its structure. The correlation between experimental binding affinity data and the values predicted by Yuel 2 is depicted in **Figure 3a**, with a Pearson correlation coefficient of 0.85. A comparison of several deep/machine learning models^11–14^ published in recent years, along with their scoring power on the same test set, is provided in **Figure 3c**. Most models achieve a Pearson correlation coefficient from 0.82 to 0.84, indicating that Yuel 2 performs at least comparably to the best among them.

**Figure 3.**
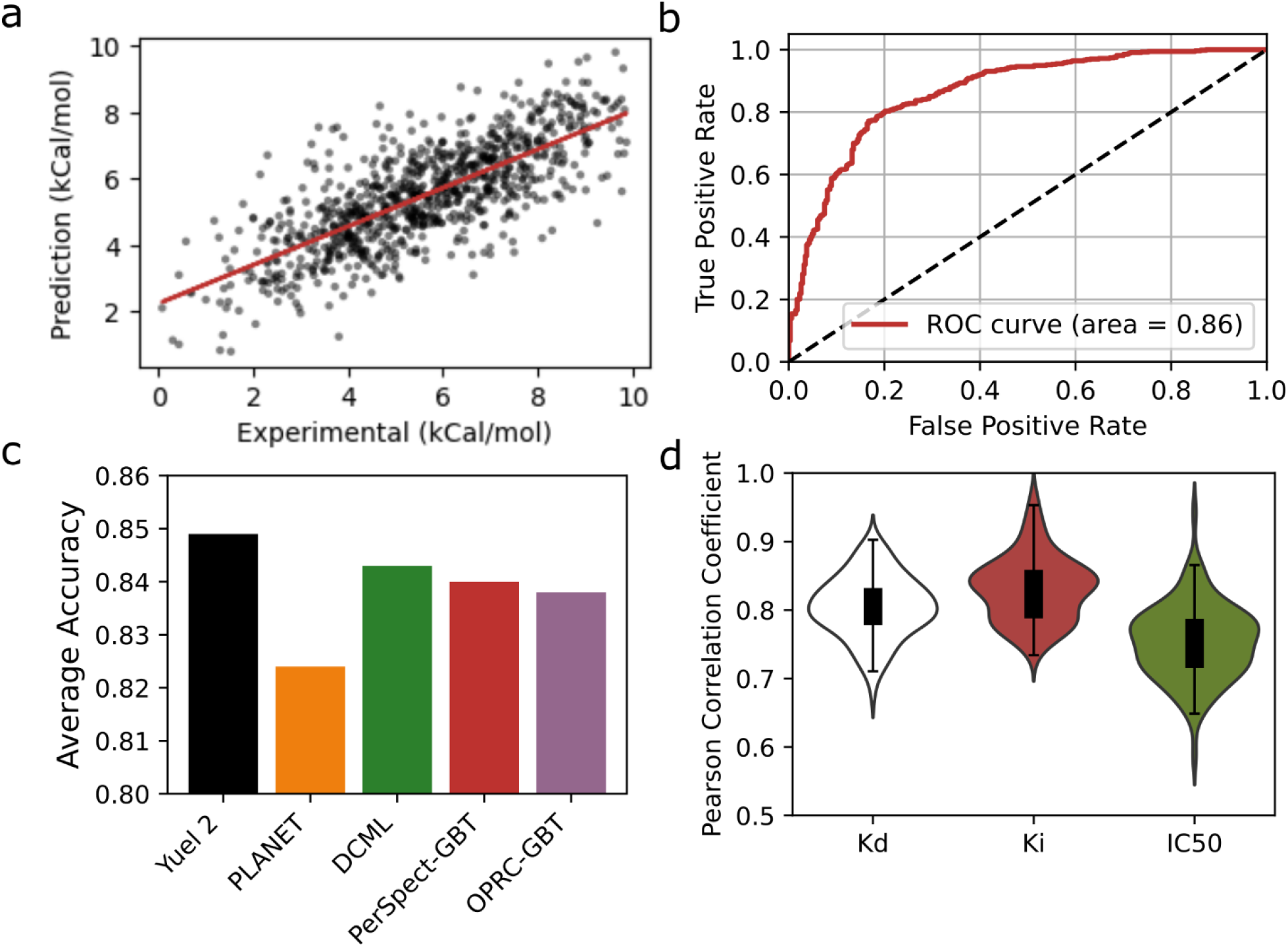
Evaluation of the scoring power and ranking power of Yuel 2. (a) The correlation between the experimental binding affinity data and the predicted binding affinity in the CASF-2016 dataset. The Pearson Correlation Coefficient is 0.85. (b) The ROC curve of the ranking power test on the CASF-2016 dataset. (c) Comparison of the average accuracy of Yuel 2 on the CASF-2016 dataset compared to the other methods. (d) Pearson Correlation Coefficient of Yuel 2 tested on the Kd, Ki, and IC50 datasets.

Unlike most other models (**Figure 3c**), which directly take 3D protein-ligand complex structures as input, Yuel 2 uses only 3D binding pocket structures and 2D ligand structures. Predicting protein-ligand binding affinity without precise binding mode information is inherently more challenging. The scoring power of deep learning models relying on crystal protein-ligand complex structures would decrease significantly if complex structures derived from molecular docking were used instead. Although the scoring power test provided by the CASF-2016 benchmark is a fundamental assessment of a protein-ligand interaction scoring function’s quality, it is not very meaningful to “predict” binding affinity based on crystal complex structures because obtaining these structures is more difficult than measuring binding affinity experimentally. Therefore, our Yuel 2 model is more robust and efficient for predicting protein-ligand binding affinity without requiring prior acquisition or generation of corresponding protein-ligand complex structures.

Another assessment provided by CASF-2016 is the ranking power test. Ranking power focuses on ranking ligands that are binding to a particular target protein by their binding affinity. This test better reflects the practical application of a scoring model. In the ranking power test, Yuel 2 achieved an average Spearman correlation coefficient of 0.682 on the 57 target proteins and an AUROC of 0.86 in the CASF-2016 test set.

Finally, we evaluate the ability of Yuel 2 to predict multiple binding metrics. PDBbind general dataset provides protein-ligand pairs characterized by three distinct metrics: Kd, Ki, and IC50. To evaluate Yuel 2, we train and test the model using the PDBbind general dataset, with an 80:20 split between the training and test sets. The results (**Figure 3d**) show that Yuel 2 achieves average Pearson correlation coefficients of 0.81 for Kd, 0.85 for Ki, and 0.75 for IC50. Overall, Yuel 2 demonstrates Pearson correlation coefficients ranging from 0.75 to 0.85, indicating a robust performance in predicting binding affinities across different metrics and datasets.

### Evaluation of the performance on the activity cliff dataset

To rigorously evaluate Yuel 2’s ability to address activity cliffs, we compile a specialized dataset from BindingDB, which is known for its extensive collection of protein-ligand binding affinities. Our goal is to assess Yuel 2’s ability to accurately predict binding affinities for ligands that are structurally similar but exhibit significantly different affinities for the same protein. We select proteins from the BindingDB dataset that have multiple associated ligands with experimentally determined binding affinities, which ensures a diverse set of ligand interactions for each protein. For each selected protein, we identify pairs of ligands with high structural similarity. Structural similarity is quantified using FP2 fingerprints and Tanimoto coefficients. For each ligand pair, we calculate the difference in their binding affinities and multiply it by their structural similarity, resulting in a metric we term the Similarity-Weighted Affinity Difference (SWAD). This metric highlights pairs where structural similarity is high, but binding affinity differences are large, making them challenging cases for prediction methods. To evaluate Yuel 2’s predictive performance, we calculate the SWAD for all ligand pairs within the same protein.

We utilize Yuel 2 to predict the binding affinities for these ligand pairs. We compute the predicted SWAD values by taking the difference in predicted binding affinities and multiplying by the structural similarity of the ligand pairs. The predicted SWAD values are compared to the true SWAD values. The Pearson correlation coefficient between the predicted and true SWAD values is calculated to quantify the predictive accuracy of Yuel 2. Our analysis yields a Pearson correlation coefficient of 0.65 (**Figure 4**), indicating Yuel 2’s ability to differentiate between ligands with similar structures that have different binding affinities. As activity cliffs represent a prominent challenge in predicting protein-ligand interactions, the ability of Yuel 2 to differentiate between small molecules on these activity cliffs represents a major milestone in our understanding of molecular interactions.

**Figure 4.**
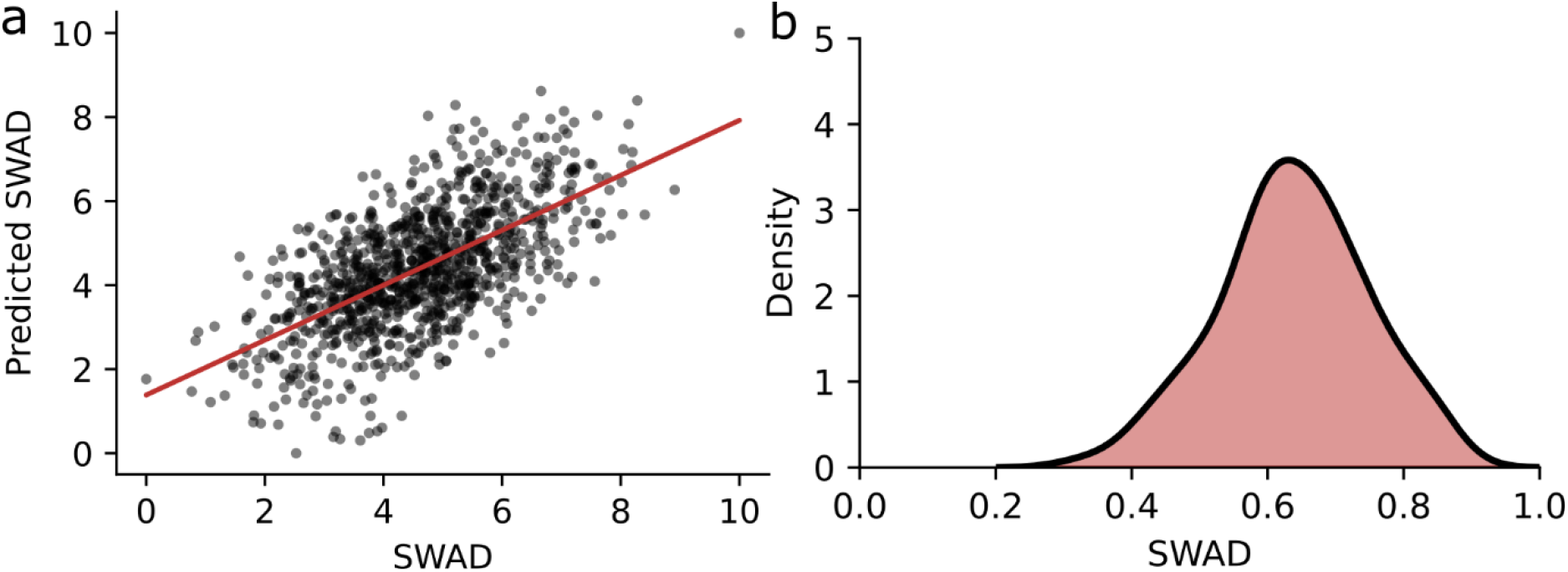
The evaluation of Yuel 2 on the activity cliffs dataset. (a) The Predicted SWAD versus the true SWAD for all pairs of ligands in the activity cliff dataset. (b) The distribution of the Pearson Correlation Coefficient of the predicted SWAD in the activity cliff dataset.

## DISCUSSION

The integration of transfer learning in Yuel 2’s approach represents a significant advancement in enhancing model performance and generalization. By employing a two-step training process, where Yuel 2 is first pre-trained on a large-scale dataset to predict docking scores and subsequently fine-tuned with a smaller, specialized dataset, we effectively leverage both extensive and focused data. The pre-training phase allows the model to learn from a diverse set of protein-ligand pairs, capturing a broad range of structural and interaction features. Fixing the parameters of Yuel-SE during the fine-tuning phase ensures that the model retains the learned structural knowledge while adapting to specific binding affinity metrics in a smaller dataset. This strategy not only improves the model’s ability to predict binding affinities accurately but also enhances its robustness across different types of data. Transfer learning thus provides a mechanism to overcome the limitations imposed by small datasets and ensures that Yuel 2 benefits from the richness of both large-scale and targeted data.

Predicting multiple metrics, including Kd, Ki, and IC50, provides a distinct advantage over models focused on a single metric. The comprehensive approach captures a broader spectrum of interactions between proteins and small molecules, offering a more nuanced understanding of their binding dynamics. While a model predicting only Kd may excel in specific scenarios, incorporating IC50 and Ki predictions enhances versatility in selecting effective drugs and a direct view of the impact of a small molecule on cell functioning. For example, IC50 represents the inhibitor concentration at which the response is reduced by half, which provides insights into inhibitory potency.

Yuel 2’s adept handling of the activity cliffs problem holds significant implications for advancing research in various domains. Activity cliffs, where small chemical modifications lead to disproportionately large changes in bioactivity, pose a formidable challenge in the design of bioactive compounds. By effectively navigating and mitigating activity cliffs, Yuel 2 contributes to the identification and optimization of compounds with subtle structural variations, ensuring a more comprehensive understanding of their bioactivity. This capability is crucial not only for drug discovery but also for other fields such as nutrition research, where fine-tuning the molecular structures of bioactive compounds, such as nutrients or nutraceuticals, can enhance their efficacy while minimizing unintended effects. This breakthrough accelerates the development of nutritionally beneficial compounds and promotes a more targeted and efficient approach to designing functional foods and supplements with optimized health benefits.

Finally, Yuel 2 can predict binding affinities using only the separate protein and ligand structures as inputs, without requiring the docking software, to obtain the complex structure. This feature significantly enhances the speed and efficiency of our approach, for it eliminates the need for time-consuming docking processes. Many existing neural network models necessitate the complex structure as an input, thereby requiring the integration of docking procedures. In contrast, our models bypass this requirement, offering a streamlined and rapid prediction workflow.

## METHODS

### The Architecture of Yuel 2

Yuel 2 comprises two main components: Yuel Structural Encoder (Yuel-SE) and Yuel Affinity Predictor (Yuel-AP). Yuel-SE is designed to extract meaningful structural features from both the protein pocket and the small molecule. During the pre-training phase, Yuel-SE is combined with Yuel-AP and trained on a large-scale dataset to predict docking scores generated by traditional docking software like MedusaDock and AutoDock Vina. This extensive pre-training allows Yuel-SE to learn the intricate structural patterns and interactions between proteins and ligands.

Yuel-SE requires two inputs to estimate interactions between a specific compound and the protein target: (i) the compound’s 2D structure (SMILES^15^ or INCHI^16^ code) and (ii) the pocket structures. We employ an in-house C++ program to extract the pocket structure from a given protein. For proteins with unknown binding pockets, we use the FPocket^17^ program to predict the position of the binding pocket. We use a graph structure to represent the pocket structure. Each node in the graph represents the “Ca” atom in a protein residue, and each edge in the graph represents the contact between two residues with features such as whether the two residues are neighboring and the distance between the two residues Ca atoms.

Given the SMILES of the compound, *Yuel-SE* first employs rdkit^18^ to represent the structure by a graph (*N, V, E*), where *N* is the number of nodes, *V* is the feature vector of each atom, and *E* is the feature vector of each bond. The feature of each atom is the concatenation of the one-hot encoding of atom type, number of bonds, bond type, mass, and charge vectors. The feature of each bond is the bond order.

The graphs of the protein and the small molecule are merged into an intact graph by adding edges between each node in the protein graph and each node in the small molecule graph (Figure 1). The merged graph is then subject to 8 graph attention layers. The node features after the graph attention layers are then subject to linear layers to predict a latent vector, which will be used as the inputs for Yuel-AP.

After pre-training, Yuel-SE’s parameters are fixed, and Yuel-AP is fine-tuned on a smaller dataset, the PDBbind general set, to predict binding affinities more accurately. Yuel-AP consists of six linear layers, taking as input the latent predicted by the Yuel-SE and directly predicting the overall binding affinity.

### Preparation of the Fine-Tuning Dataset

The PDBbind “general set” is a comprehensive collection of protein-ligand complexes with available 3D structures from the Protein Data Bank (PDB) and corresponding experimental binding affinity data (i.e., Kd, Ki, and IC50) curated from the literature. This dataset has been extensively used for developing deep learning and machine learning models aimed at predicting protein-ligand binding affinity. For our study, we used the PDBbind general set (v.2020), comprised of a total of 19,443 protein-ligand complexes, as the initial pool for training Yuel 2.

The PDBbind general set provides the protein PDB structure file, small molecule SDF and Mol2 files, as well as the PDB structure file of the pocket. We use an in-house C++ program to process the pocket PDB structure to convert it to a graph file. We use RDKit to process the small molecule Mol2 file and remove those that could not pass the sanitize operation, resulting in 19,386 successfully processed protein-ligand complexes from the PDBbind general set (v.2020).

### Pre-training and Fine-Tuning Dataset

To compile a large-scale dataset for pre-training, we randomly selected 500 PDB IDs from the PDBbind general set. For each selected PDB ID, we perform MedusaDock docking attempts and AutoDock vina docking attempts to obtain the MedusaScore energies and AutoDock scores. The docked file and the docking scores are stored in a Sqlite 3 database file. This approach yielded a total of 500 × 500 = 250,000 protein-ligand pairs, creating a diverse and extensive dataset for pre-training. The PDBbind general set is used as the fine-tuning dataset. We split it to the training and test sets with a ratio of 8:2.

Finally, we use the CASF-2016 dataset for the comparison of Yuel 2 to other methods. To ensure the reliability of our model, we excluded the 285 protein-ligand complexes in the CASF-2016 test set from our training set.

### Activity Cliffs Dataset

To evaluate Yuel 2’s performance in addressing activity cliffs, where structurally similar compounds exhibit markedly different binding affinities for a single protein target, we propose the following approach to compile a dataset. We begin with the BindingDB dataset, which provides a comprehensive repository of protein-ligand binding affinities. The process involves several steps to isolate relevant examples of activity cliffs. First, we identify proteins with a significant number of binding affinities reported for multiple ligands in the BindingDB dataset. This ensures that the dataset will contain a sufficient variety of compounds for each protein target. For each selected protein, we extract a subset of ligands that have similar chemical structures. This is achieved by clustering compounds based on their molecular fingerprints or using chemical similarity metrics. Within each cluster of structurally similar compounds, we identify cases where there is a substantial variation in binding affinities. These instances are characterized by a high discrepancy in affinity values despite the close structural resemblance among the compounds. We then construct a dataset that includes these pairs of similar compounds with divergent binding affinities. Each entry should contain the protein target, structurally similar compounds, and their corresponding binding affinities. We format the binding affinity information alongside the molecular representations of the ligands and the protein structure. Ensuring that the dataset is split into training and validation subsets, we may properly assess Yuel 2’s ability to accurately predict binding affinities and handle activity cliffs.

### Model evaluation

In the context of computational chemistry and drug discovery, “scoring power” refers to the ability of a scoring model to predict a binding score that exhibits a linear correlation with experimental protein−ligand binding data, based on valid inputs. In our study, we evaluate the scoring power of Yuel 2 using the CASF-2016 benchmark. CASF-2016 is a widely recognized benchmark for assessing scoring functions, including various machine learning models (see Table 1 for examples). The test set in CASF-2016 comprises 285 diverse protein−ligand complexes with high-quality crystal structures and experimental binding data.

In alignment with the CASF-2016 protocol, our evaluation metrics include the Pearson correlation coefficient between experimental binding data and predicted values. Another related evaluation is the ranking power test. “Ranking power” assesses the ability of a scoring model to correctly rank known ligands of a common target protein. Unlike scoring power, ranking power measures how well the model ranks ligands based on their binding data, rather than expecting a linear correlation between predicted binding data and actual values. For each target protein, we computed the Spearman correlation coefficient between experimental binding data and predicted values. The overall ranking power was determined by averaging the Spearman correlation coefficient values across all 57 target proteins.

## DATA AVAILABILITY

Source codes and test data are deposited at: *https://bitbucket.org/dokhlab/yuel2*.

## ACKNOWLEDGEMENTS

We acknowledge support from the National Institutes for Health (R35 GM134864), the National Science Foundation (2040667), and the Passan Foundation. This project was also supported by the Penn State College of Medicine’s Artificial Intelligence and Biomedical Informatics Program. We appreciate Congzhou Sha and Alicia Xie for their enthusiastic proofreading of the manuscript.

## SUPPORTING INFORMATION AVAILABLE

The supporting information provides extra testing results of Yuel 2.

