## Supplemental Information for "Leveraging Transfer Learning for Predicting Protein-Small Molecule Interactions"

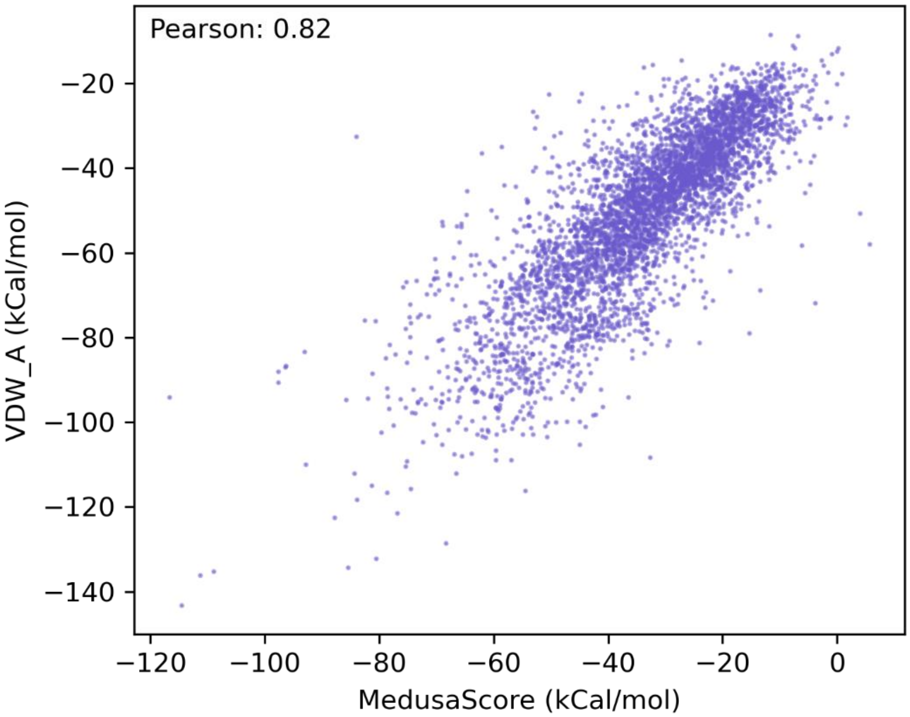


**Figure S1. VDW_A has a high relationship with MedusaScore**

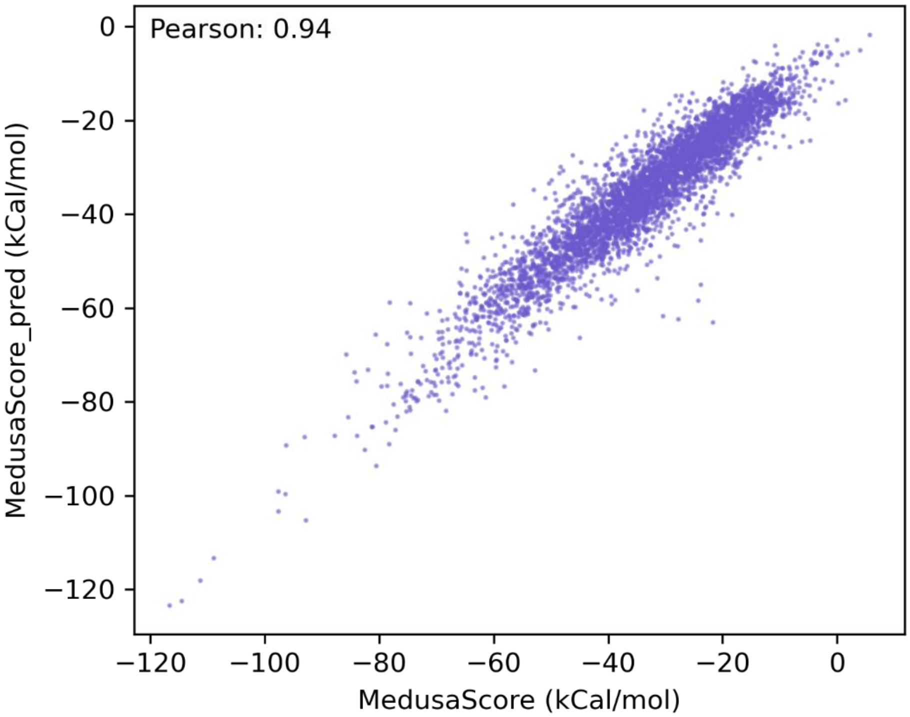


**Figure S2. Yuel 2 can predict the total MedusaScore with a Pearson correlation coefficient of 0.94.**

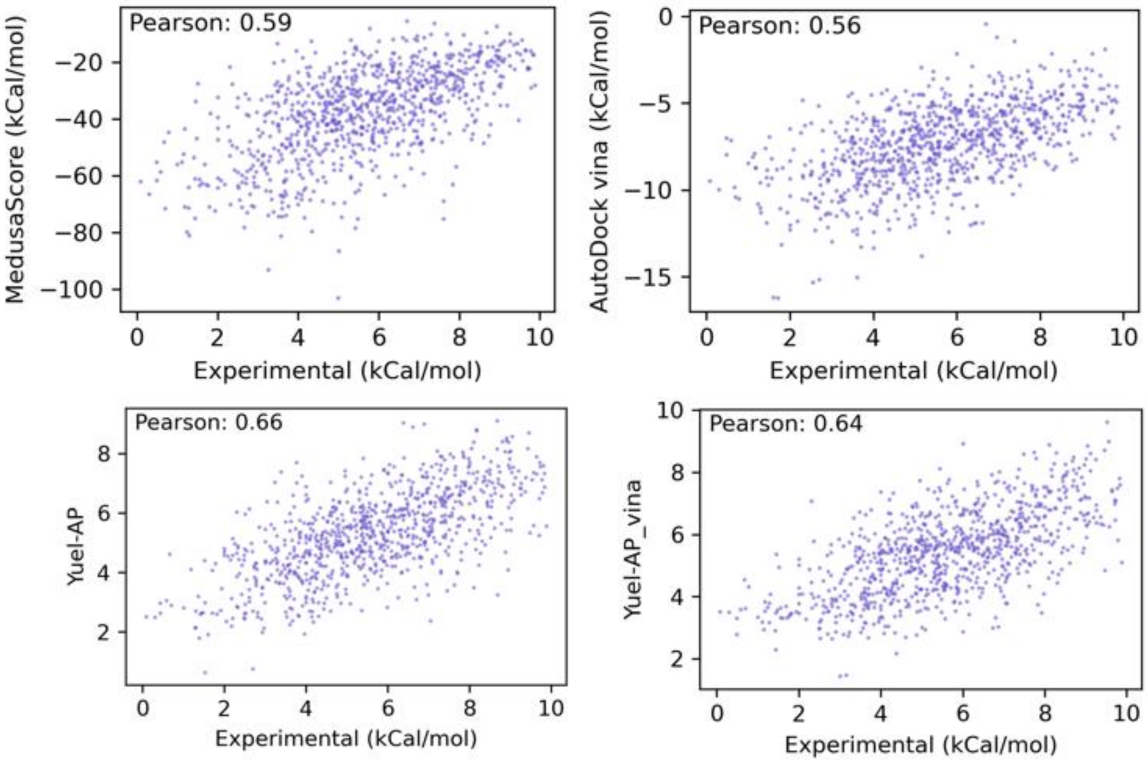


**Figure S3. Yuel 2 improves the performance of MedusaDock and AutoDock vina.**
